# Development of Functional Connectome Gradients during Childhood and Adolescence

**DOI:** 10.1101/2021.08.08.455594

**Authors:** Yunman Xia, Mingrui Xia, Jin Liu, Xuhong Liao, Tianyuan Lei, Xinyu Liang, Tengda Zhao, Ziyi Shi, Lianglong Sun, Xiaodan Chen, Weiwei Men, Yanpei Wang, Zhiying Pan, Jie Luo, Siya Peng, Menglu Chen, Lei Hao, Shuping Tan, Jiahong Gao, Shaozheng Qin, Gaolang Gong, Sha Tao, Qi Dong, Yong He

## Abstract

Connectome mapping studies have documented a principal primary-to-transmodal gradient in the adult brain network, capturing a functional spectrum which ranges from perception and action to abstract cognition. However, how this gradient pattern develops and whether its development is linked to cognitive growth, topological reorganization, and gene expression profiles remain largely unknown. Using longitudinal resting-state functional magnetic resonance imaging data from 305 children (ages 6-14), we describe substantial changes in the primary-to-transmodal gradient between childhood and adolescence, including emergence as the principal gradient, expansion of global topography, and focal tuning in primary and default-mode regions. These gradient changes are mediated by developmental changes in network integration and segregation, and are associated with abstract processing functions such as working memory and expression levels of calcium ion regulated exocytosis, synaptic transmission, and axon and synapse part related genes. Our findings have implications for understanding connectome maturation principles in normal development and developmental disorders.

**Teaser:** Our study reported the maturation of the core connectome gradient and its association with cognitions and genes expression.

## Introduction

Hierarchy is a fundamental organizational principle in the human brain, and promotes the encoding of information from low-level sensation into high-order cognitive processing (1, 2). Classical neuroanatomy studies based on postmortem brains have documented hierarchical mechanisms in the organization of cortical microstructures, such as the distribution of neuronal cells (cytoarchitecture) and myelinated nerve fibers (myeloarchitecture) (2–4). With advances in non-invasive neuroimaging techniques (5) making the decomposition of connectivity data from the human brain possible (6), researchers have demonstrated the presence of a principal connectivity gradient in the macroscale organization of functional brain networks in healthy adults. This principal connectivity gradient spans the primary sensory network and the transmodal regions of the default-mode network (DMN) (6, 7). The connectome hierarchy reflected in this gradient captures a functional spectrum from perception and action to increasingly abstract cognitive domains, suggesting that it plays a crucial role in supporting complex cognitive functions and behaviors (1, 6).

Despite such importance, developmental changes in the core primary-to-transmodal gradient remain understudied. A previous study reported that this gradient pattern does not present in the neonatal connectome, reflecting an immature differentiation between sensory and transmodal regions (8). Although a recent study has found the presence of the principal gradient of connectivity in adolescence (9), how this gradient pattern develops in children and whether its development is linked to cognitive growth, topological reorganization, and gene expression profiles remain poorly understood. Thus, the maturation process of the principle connectome gradient and its cognitive implication and biological basis remain to be established.

Childhood and adolescence are critical periods for the development of a wide range of cognitive and behavioral capabilities, including fine motor skills, abstract thinking, and complex semantic cognition (10). In particular, according to Piaget’s theory of cognitive development, the transition from the concrete operational stage (ages 7-11) to the formal operational stage (ages 12 and up) happens between middle childhood and early adolescence. In the course of this transition, cognitive development in children gradually shifts from concrete to abstract and logical thinking (11). During this period, the functional brain connectome underlying cognitive growth undergoes rapid and substantial reconfiguration and refinement (12–14). Functional connectivity studies have revealed a significant increase in anterior-posterior connections within the DMN (15, 16), and further show that the DMN develops from a grouping of sparsely functionally connected regions to a cohesive and interconnected network (15, 17). Studies based on graph-theoretical analysis suggest that substantial network (e.g., DMN) balancing changes, including both regional segregation of anatomically neighboring areas, and global integration between distant regions (18–20) also occur between middle childhood and early adolescence. Moreover, several regions of the DMN (e.g., the medial prefrontal and parietal cortices) progressively play central roles in the whole-brain network as hubs, linking functional communities and promoting efficient communication (21, 22). Results from these studies imply that the DMN undergoes a transition from underdeveloped to matured during this age period. At the same time, the DMN also plays an important role in the adult primary-to-transmodal gradient, serving as an anchor in the gradient axis. Thus, we speculated that maturation of the DMN between middle childhood and early adolescence may drive a dramatic development in the primary-to-transmodal gradient. However, how this connectome gradient pattern emerges and develops during childhood and adolescence remains largely unknown.

Nonetheless, it has been well established that structural and functional development of the brain is precisely regulated by genetic factors (23, 24). For example, the Catechol-O-metyltransferase (COMT) genotype, which is related to synaptic levels of dopamine, affects prefrontal white matter pathways in the child brain (25). The brain-derived neurotrophic factor (BDNF) gene, known to regulate neurogenesis and synaptic formation, affects functional connectivity within the DMN in children (26). However, it is extremely difficult to directly measure the expression levels of these neurodevelopment-related genes *in vivo*. Recently, researchers have been able to bridge the gap between macroscale functional brain organization and microscale biological processes through the combined investigation of connectomes derived from neuroimaging and gene expression profiles sampled from the postmortem brain (27–30). Several connectome-transcriptome association studies have demonstrated a nexus between functional connectivity in the brain and the expression levels of genes enriched for ion channel activity, synaptic function, and aerobic glycolysis (29, 31). Thus, we speculate that the developmental changes in the principal primary-to-transmodal gradient during childhood and adolescence will be influenced by the expression levels of neurodevelopment-related genes.

To address these issues, we collected a large dataset of longitudinal resting-state functional MRI (rs-fMRI) scans from a cohort of 305 typically developing children (ages 6-14, 491 scans in total). Using a previously described connectome gradient decomposition technique (6), we first identified the principal connectivity gradients in the functional connectome and then investigated their development during childhood and adolescence. Further, we conducted a connectome-transcriptome association analysis to investigate transcriptional profiles related to the development of connectome hierarchy by using gene expression data (obtained from postmortem brain samples) from the Allen Institute for Brain Science (27). In undertaking these investigations, we sought to test two hypotheses: (*i*) that the connectome gradient spanning the primary and transmodal regions is present during childhood and continues to develop throughout childhood and adolescence; and (*ii*) that developmental changes in the principal primary-to-transmodal gradient are linked to the expression profile of specific genes, such as synaptic-related genes, that are associated with brain development.

## Results

To test our hypotheses, we undertook a longitudinal study on a cohort of 305 typically developing children (ages 6-14) (“Discovery Dataset”). Each child underwent up to three separate rs-fMRI scans taken at different points in time, resulting in a total of 491 longitudinal scans. An independent dataset (“Replication Dataset”) from the enhanced Nathan Kline Institute-Rockland Sample (NKI-RS) was used to validate our main results. This Replication Dataset consisted of longitudinal data from a cohort of 254 typically developing children (ages 7-18, N = 390 scans). Rs-fMRI scans from a sample of 61 healthy young adults (ages 18-28) were also taken, using the same scan parameters as Discovery Dataset, to construct a reference dataset of the mature human brain and allow for cross-sectional comparisons. All MRI scans were subjected to a strict quality control selection process before inclusion in this study. Details on participation inclusion, data collection, and quality control are provided in the *SI Appendix*, Section 1-3.

We began our analysis by examining functional connectivity gradients (6) in healthy young adults with mature connectomes (N = 61). Specifically, we first constructed an individual functional connectivity matrix at the voxel level for each participant and obtained a group averaged functional network. Then, we used the diffusion map embedding method to decompose the group-level network into multiple gradients representing the connectivity variance in the functional connectome (*SI Appendix*, Section 4, Fig. S1). These gradients were mapped onto the cortical surface to define the macroscale pattern, and each brain region (i.e., voxel) was assigned a gradient score. The gradient score reflects the relative position of brain regions along the gradient axes, which is determined by the difference in connectivity patterns of brain regions (6). We found that the principal gradient, which accounted for the greatest variance in the functional connectome (29.37%, Fig. 1A), as derived from our dataset, was anchored at one end by the primary sensory network (which includes the visual and sensorimotor regions) and at the other by the transmodal regions of the DMN (Fig. 1B, *top*). The second most significant gradient accounted for 22.82% of connectome variance in our dataset (Fig. 1A), with a gradual axis defined by the sensorimotor network at one end and the visual network at the other (Fig. 1B, *bottom*). These results replicate prior findings with respect to connectome hierarchy in healthy adults (Fig. 1C) (6). Given that these two gradients (i.e., the primary-to-transmodal gradient and the sensorimotor-to-visual gradient) account for the greater part of connectome variance (52.19% in total), and given their association with neuronal microstructure and relevance to cognitive functions (8, 32, 33), the present study focuses primarily on the developmental process of these two gradients.

**Fig. 1.**
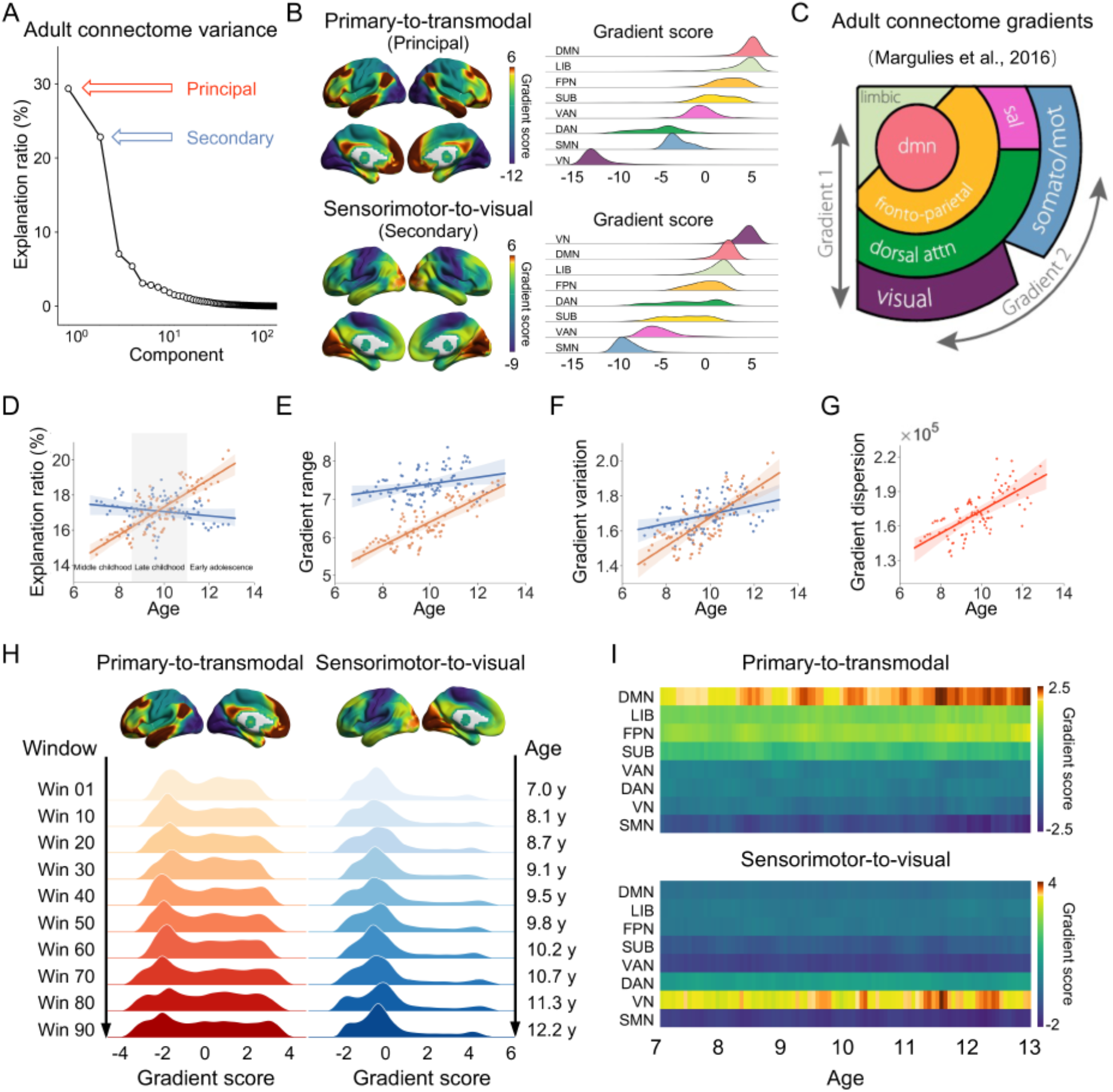
Connectome gradients in adults and children. (**A**) The explanation ratio of connectome gradients in adults. The principal gradient accounts for 29.37% of the connectivity variance in the functional connectome, and the second most significant gradient accounts for 22.82%. (**B**) The core connectome gradients in adults. The principal gradient is anchored at one end by the primary sensory network and at the other end by the transmodal regions of the DMN (*left top*). The second core gradient is defined by the SMN at one end and the visual cortex at the other (*left bottom*). System-level distributions of the two gradients (**B**, *right*) are highly consistent with findings in prior studies (**C**) (6). Global measures used to quantify gradient properties of the two connectome gradients across age windows were the explanation ratio (**D**), gradient range (**E**), gradient variation (**F**), and gradient dispersion (**G**). Gradient dispersion and global measures in the primary-to-transmodal gradient increase continuously with age, while those in the sensorimotor-to-visual gradient remain relatively stable. Gradient global measures of the primary-to-transmodal gradient and gradient dispersion values are indicated in orange and red respectively, while gradient global measures of the sensorimotor-to-visual gradient are indicated in blue. (**H**) Global histograms showing the gradient scores for each of the two gradients across representative age windows. The histograms of the primary-to-transmodal gradient (*left*) show a trend of increasing spread with age, while those of the sensorimotor-to-visual gradient (*right*) remain relatively constant. (**I**) Mean gradient scores in different functional systems across age windows. Primary-to-transmodal gradient scores in the SMN decreased with age, while those in the DMN increased with age. Surface rendering was generated using BrainNet Viewer (www.nitrc.org/projects/bnv/) (78). VN, visual network; SMN, sensorimotor network; DAN, dorsal attention network; VAN, ventral attention network; SUB, sub-cortical regions; LIB, limbic network; FPN, frontoparietal network; DMN, default-mode network.

### The Primary-to-Transmodal Gradient Exhibits Both Global and Regional Development and Accounts for an Increasing Proportion of Connectivity Variance

To delineate the development of connectome hierarchy during childhood and adolescence, we conducted both a qualitative cross-participant sliding window analysis and a quantitative mixed linear model analysis on the 491 child brain scans in the Discovery Dataset (for details, see *SI Appendix*, Section 4-5). Prior to undertaking these analyses, we used the diffusion map embedding method (6) to decompose the child brain functional connectomes derived from the scans into connectivity gradients, and further applied Procrustes rotation approaches to align the connectivity gradients from each scan to a group-based, iterative gradient template. These procedures ensure that, throughout our study, analyses are being conducted on the same gradient(s) across all scans in the dataset. Notably, the two core gradients (i.e., the primary-to-transmodal gradient and the sensorimotor-to-visual gradient), as observed in the empirically obtained gradient distribution patterns, accounted for significantly greater connectivity variance as compared to corresponding gradients in the rewired null model for each child scan (all *P* < 0.033, *SI Appendix*, Section 4).

#### Cross-participant sliding window analyses

For this analysis, we divided the child brain scans in the Discovery Dataset into different age windows in ascending order of age (*SI Appendix*, Fig. S2). Each window consisted of 30 scans and had a step size of five (i.e., window 1 contained scans 1-30, window 2 contained scans 6-35, …, window 93 contained scans 461-491). This processing generated 93 overlapping subgroups of brain scans, with the average age of children in each subgroup ranging from 7.0 to 12.9 years. Within each window, we averaged the aligned gradients across all scans. We investigated the changes in connectome gradients with age using four global measures, being the explanation ratio (6), the gradient range (32), the gradient variation, and the gradient dispersion (7). For a given gradient, its explanation ratio represents the percentage of connectivity variance accounted for by that gradient. A greater explanation ratio suggests that the embedding axis of the given gradient captures a more dominant organization of the functional connectome. The gradient range captures the difference between the greatest positive and negative values of brain voxels in a given gradient. A larger range indicates a greater difference in the encoded connectivity pattern between the regions localized at the gradient ends. The gradient variation represents the standard deviation of the gradient scores across the whole brain. A greater variation reflects a higher heterogeneity in the connectivity structure across regions. The gradient dispersion captures the summed square of the Euclidean distance from each brain voxel to the centroid of the gradient space. A greater dispersion means a more discrete distribution of brain regions in the gradient space determined by the first two gradients (7).

Explanation ratios obtained from our dataset showed that, during the years spanning childhood and adolescence, the primary-to-transmodal gradient transitions from being the second most significant connectivity gradient into becoming the principal gradient in the functional connectome (Fig. 1D). Specifically, in middle childhood (ages 7-8.5), the primary-to-transmodal gradient accounts for the second-largest percentage of connectivity variance in the functional connectome, ranking behind the sensorimotor-to-visual gradient (*t* = -2.19, *P* = 0.031, paired *t*-test on explanation ratio), which is the principal connectivity gradient during this stage of development. These middle childhood explanation ratios match those observed in neonatal connectome gradients (8). In late childhood (ages 8.5-11), the primary-to-transmodal and sensorimotor-to-visual gradients exhibit relatively close explanation ratios (*t* = -0.22, *P* = 0.828, paired *t*-test on explanation ratio). By early adolescence (ages 11-13), the primary-to-transmodal gradient begins to transition into the principal connectivity gradient (*t* = 2.82, *P* = 0.006, paired *t*-test on explanation ratio), indicating that this is a critical phase for the emergence of a typical adult-like connectome gradient. The gradient range and variation of the primary-to-transmodal gradient and the gradient dispersion also exhibit age-related increases, suggesting gradually maturing global topographic profiles in children (Fig. 1E-1G). This is further confirmed by observations in our study that the global histograms exhibit greater spread in gradient scores with age (Fig. 1H). According to Yeo’s functional parcellations (34), gradient scores in the DMN gradually increased with age, while those in the sensorimotor network (SMN) gradually decreased with age (Fig. 1I, *top*), indicating an enlarged dissimilarity in the embedding functional connectivity pattern between the DMN and SMN regions. In contrast, all global and regional measures in the sensorimotor-to-visual gradient were relatively stable across the different age windows (Fig. 1D-1F, 1H and 1I *bottom*). Overall, these qualitative analyses suggest that the primary-to-transmodal gradient emerges as the principal gradient in the connectome around early adolescence and continues to account for an increasing proportion of connectivity variance as the functional connectome develops towards a more adult-like hierarchical organization.

#### Mixed linear model analyses

Next, we used a mixed linear model to conduct a quantitative analysis of the scans from the Discovery Dataset (Fig. 2A) to examine the longitudinal changes in the two connectome gradients (*SI Appendix*, Section 5). In this model, we assessed the effect of age on connectome gradient measures, with gender and head-motion parameters included as co-variates. We observed significant age-associated increases between childhood and adolescence in the following: (*i*) all three global measures of the primary-to-transmodal gradient (explanation ratio: *t* = 5.22, *P <* 0.001, Fig. 2B; gradient range: *t* = 5.13, *P <* 0.001, Fig. 2C; gradient variation: *t* = 5.21, *P <* 0.001, Fig. 2D); (*ii*) the difference in the explained ratio of the two core gradients (*t* = 3.55, *P <* 0.001, Fig. 2E); and (*iii*) gradient dispersion of the two core gradients (*t* = 3.73, *P* < 0.001, Fig. 2F). Voxel-wise statistical analysis revealed that age-associated increases in gradient scores were mainly concentrated in the high-order DMN regions, such as the medial and lateral prefrontal and parietal cortices, and the lateral temporal cortex, while age-associated decreases mapped to primary regions, such as the sensorimotor, auditory, and visual regions (voxel-level *P <* 0.001, Gaussian random field cluster-level corrected *P <* 0.05, Fig. 2G and Table S1). These age effects remained significant after controlling for the confounding effects of the sensorimotor-to-visual gradient (*SI Appendix*, Fig. S3). For the sensorimotor-to-visual gradient, we did not observe age effects in any of the global measures (*P* > 0.05 for all three global measures: explanation ratio, gradient range, and gradient variation). Regionally, age-associated changes in gradient scores were observed in a few sites, including the middle occipital lobe, the inferior parietal gyrus, and the superior temporal gyrus (voxel-level *P <* 0.001, Gaussian random field cluster-level corrected *P <* 0.05) (*SI Appendix*, Fig. S4 and Table S1).

**Fig. 2.**
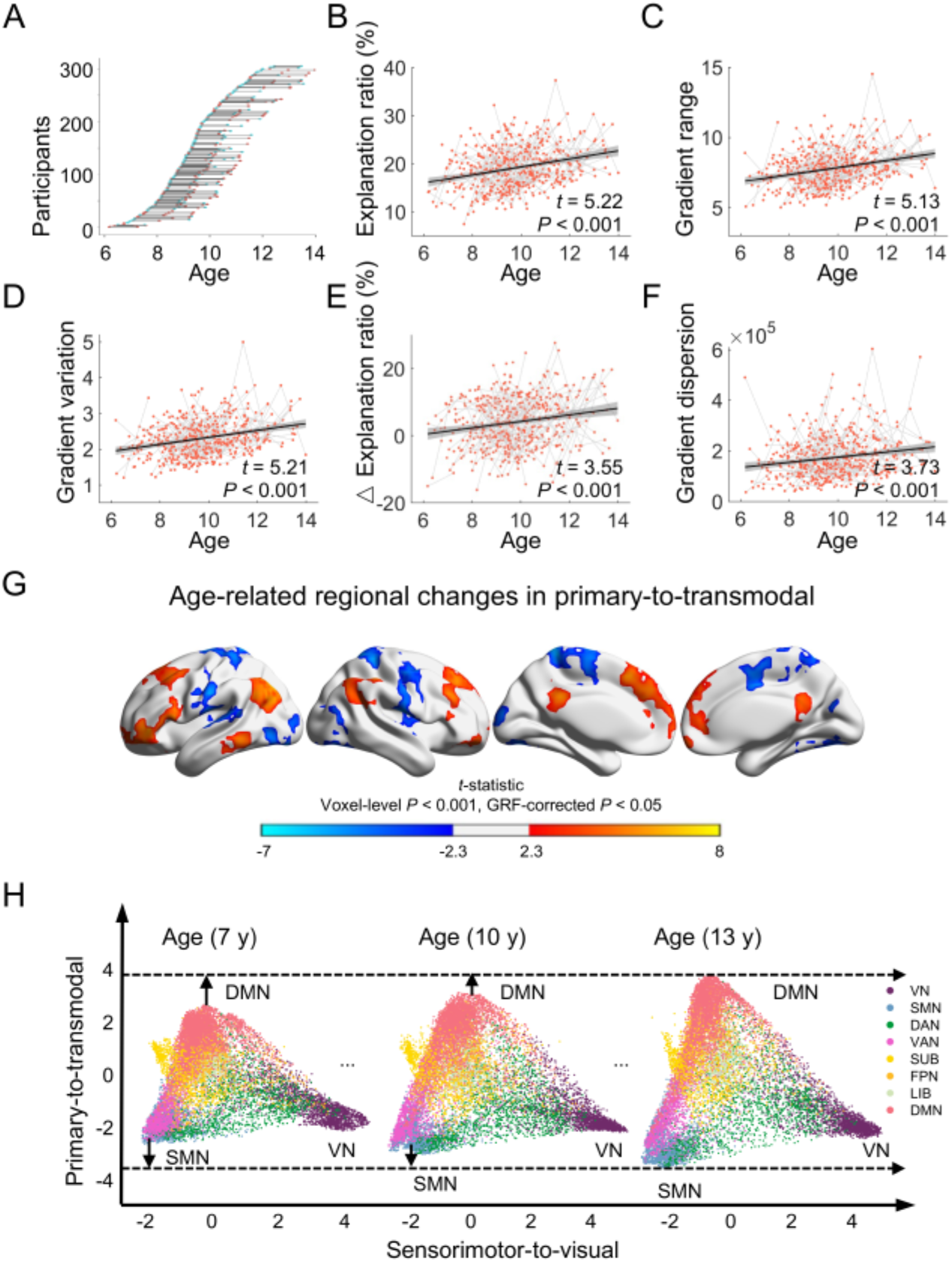
Longitudinal changes in the primary-to-transmodal gradient in children. (**A**)Age of subjects when scans were taken. Each dot represents one child brain scan. The blue dots represent the scans from males, while the red dots represent the scans from females. Each line represents the time interval between scans of the same subject. Significant increases with age were seen in all three global measures of the primary-to-transmodal gradient, including the explanation ratio (**B**), gradient range (**C**), and gradient variation (**D**), the difference in the explanation ratio of the two core gradients (**E**), and gradient dispersion (**F**). (**G**) At the regional level, gradient scores that increased with age were mostly found within the DMN (shown in warm colors), while gradient scores in the VN and SMN decreased with age (shown in cool colors) (voxel-level *P* < 0.001, Gaussian random field cluster-level corrected *P* < 0.05). For specific sites see *SI Appendix, Table S1*. (**H**) Scatter plots of voxel-wise gradient scores for each of the two connectome gradients at three representative ages (7 years old, 10 years old, and 13 years old). In the primary-to-transmodal gradient (*y-axis*), larger differences are observed between gradient scores in the primary regions and in the DMN. In the sensorimotor-to-visual gradient (*x-axis*), regional gradient scores are relatively fixed across the three ages. Each dot represents a voxel and is colored according to its functional system assignment (34).

To better visualize the longitudinal changes in the connectome gradients, we established a co-ordinate system, with gradient scores for the primary-to-transmodal gradient represented in the *y-axis*, and gradient scores for the sensorimotor-to-visual gradient represented in the *x-axis* (Fig. 2H). Scatter plots of three representative ages (7 years old, 10 years old, and 13 years old) were illustrated, representing the gradient scores for each voxel across the major functional systems in the connectome from middle childhood to late childhood and through to early adolescence. The synoptic mapping of these connectome gradients shows an expansion in the gradient range of the primary-to-transmodal component, which is observed in both the upper extreme (found mostly in the DMN) and lower extreme (found mostly in the primary sensory network) of the gradient scores. This indicates a trend towards differentiated connection profiles between the primary sensory network and the DMN (the developmental trajectory of these gradient hierarchies across different ages is available in the *SI Appendix*, Movie S1).

Using the Replication Dataset (ages 7-18) from the NKI-RS, we validated our main findings on the longitudinal development of connectome gradients during childhood and adolescence (*SI Appendix*, Section 5). Significant age-related changes were observed in the global measures of the primary-to-transmodal gradient (all *P* < 0.03), but not in measures of the sensorimotor-to-visual gradient (*SI Appendix*, Fig. S5). Age-related changes in regional gradient scores of the primary-to-transmodal were mainly observed in the DMN regions (*SI Appendix*, Fig. S5). These results are consistent with those obtained from our Discovery Dataset. Together, these findings from two independent datasets provide replicable evidence of the age-associated changes in core connectome gradients during childhood and adolescence.

We also performed a cross-sectional comparison of the primary-to-transmodal and sensorimotor-to-visual gradients between child and adult brains (*SI Appendix*, Section 5, Fig. S6). Specifically, all three global measures (i.e., explanation ratio, gradient range, and gradient variation) of the primary-to-transmodal gradient, as well as the gradient dispersion showed smaller values in children than adults (all *P* < 0.005). Moreover, there was a significant difference in regional gradient scores between the child and adult brains (*SI Appendix*, Fig. S6). This suggests that maturation of the core primary-to-transmodal gradient from childhood to adulthood proceeds in an ongoing and progressive manner.

### Association Between the Primary-to-Transmodal Gradient and Cognitive Functions

To examine the cognitive implications of primary-to-transmodal gradient development, we performed the following two analyses (*SI Appendix*, Section 6). First, we conducted a meta-analysis to identify cognitive terms associated with the longitudinal developmental map of the primary-to-transmodal gradient using Neurosynth (35). In this analysis, we first constructed a spatial map showing regions with significant changes in gradient score during development. We then examined the spatial correlation between the maps of regions with significant gradient score changes and meta-analytic activation maps of multiple cognitive tasks (NeuroSynth datasets). A permutation test was used to estimate the significance of the correlation coefficients, and spatial autocorrelation was corrected for using generative modeling (36). For the primary-to-transmodal gradient, brain regions with age-associated increases in gradient score were significantly associated with DMN-related cognitive functions, such as memory retrieval, judgment, semantic cognition and reasoning (Fig. 3A, *left*, FDR-corrected *q* < 0.05); brain regions with age-associated decreases in gradient score were significantly involved in somatosensory, sensorimotor and primary motor functions (Fig. 3A, *right*, FDR-corrected *q* < 0.05). For the sensorimotor-to-visual gradient, brain regions with age-associated changes in gradient score were significantly correlated with those functions usually activated when performing visual and mathematic-related tasks (*SI Appendix*, Fig. S4, FDR-corrected *q* < 0.05).

**Fig. 3.**
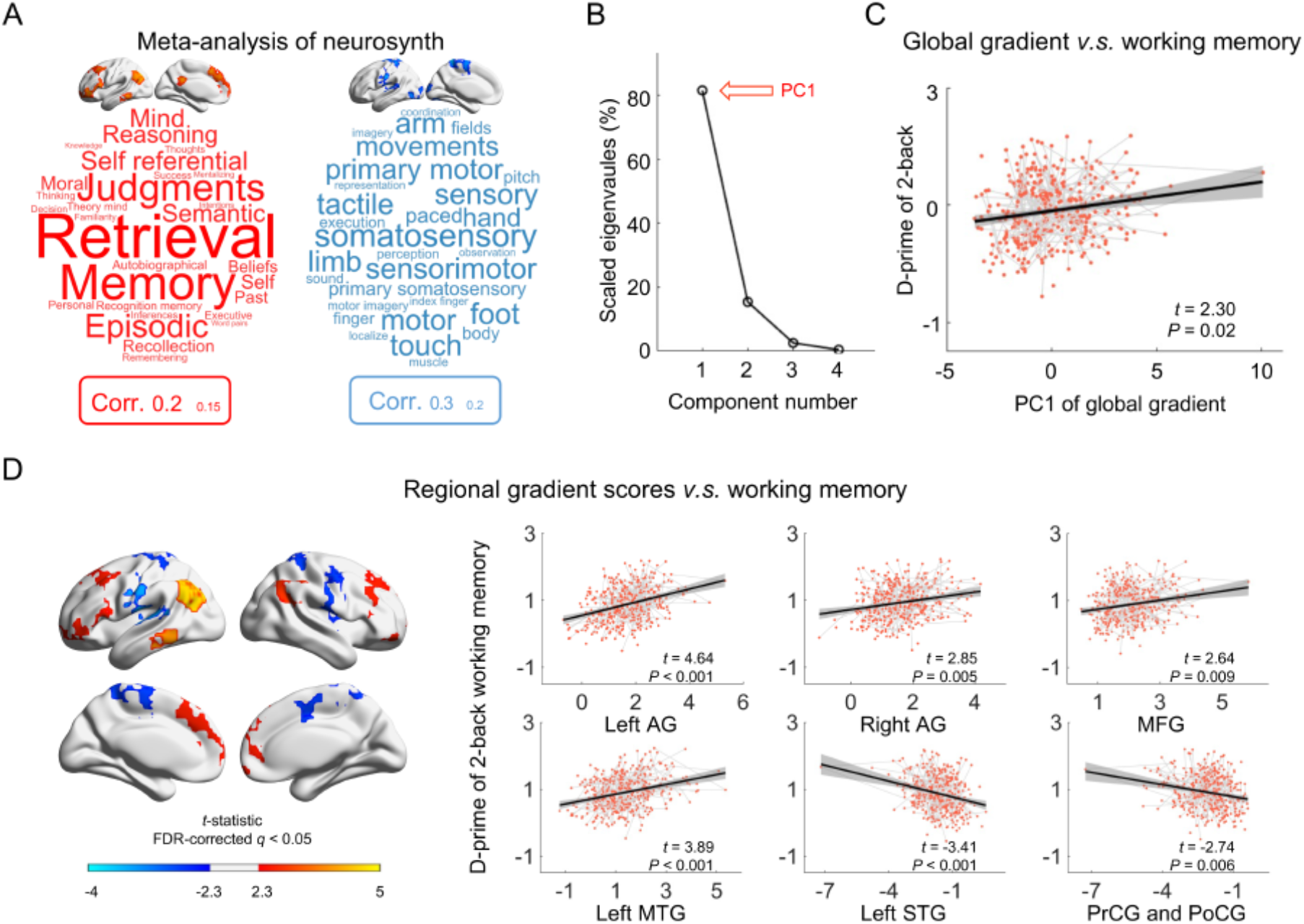
Association between maturation of the primary-to-transmodal gradient and cognitive functions. (**A**) In the primary-to-transmodal gradient, regions with age-associated increases in gradient score are associated with DMN-related functions (shown in red, *top*), e.g., memory retrieval, judgment, semantic cognition, and reasoning; regions with age-associated decreases were mainly involved in somatosensory, sensorimotor, and primary motor functions (shown in blue, *bottom*) (FDR-corrected *q* < 0.05). The font size of a given cognitive topic term represents the correlation between the age-associated *t*-statistical map and meta-analytic map for that term generated by Neurosynth. The boxes below the brain maps indicate the correspondence between font size and the correlation coefficient. (**B**) A principal component analysis (PCA) to identify the common factor of four global gradient measures exhibiting age-associated change. The first principal component (PC1) accounts for 81.55% of the variance in global gradient topography. (**C**) Association between global measures of the primary-to-transmodal gradient and performance in the *N*-back working memory task (2-back d-prime). Individual working memory capacity was positively correlated with PC1. (**D**) Association between regional gradient scores of the primary-to-transmodal and individual working memory. For each regional gradient score exhibiting age-associated changes, mean gradient scores that positively correlated with performance in the 2-back working memory task were mostly found within the DMN (shown in warm colors), while age-related changes in gradient score in the VN and SMN were negatively associated with performance in the 2-back task (shown in cool colors, FDR-corrected *q* < 0.05). AG, angular gyrus; MFG, middle frontal gyrus; MTG, middle temporal gyrus; STG, superior temporal gyrus; PrCG, precentral gyrus; PoCG, postcentral gyrus.

Second, to establish a direct link between the age-associated changes in the primary-to-transmodal and the development of specific cognitive functions, we used a mixed linear model to examine the association between the gradient measures and cognitive performance in children. Here, two cognitive dimensions were considered: (*i*) working memory, which is important for multiple higher-level abstract cognitive functions (37); and (*ii*) attentional ability, which reflects the allocation of cognitive processing resources to sensory signals (38). Given the high correlation between any pair of the four global gradient measures (i.e., the explanation ratio, gradient range, and gradient variation of the primary-to-transmodal gradient, as well as the gradient dispersion of the two gradients), we performed a principal component analysis (PCA) to reduce the dimensionality of this variable. The first PCA component (PC1, accounting for 81.55% of variance, Fig. 3B) was used to represent the common factor of global gradient measures. Controlling for gender, head-motion parameters, and baseline levels (0-back d-prime), we found that the d-prime of 2-back working memory task showed significant positive association with PC1 of the global gradient measures (*t* = 2.30, *P* = 0.02, Fig. 3C). We also analyzed the correlation between each individual global gradient measure and working memory performance (*SI Appendix*, Fig. S7). Regionally, better working memory performance was associated with the higher gradient scores in the DMN regions and lower gradient scores in sensorimotor cortex (Fig. 3D and Table S2, FDR-corrected *q* < 0.05). In contrast, we observed no association between attentional ability and global gradient measures. These results indicate that changes in primary-to-transmodal gradient during brain maturation are related primarily to the development of abstract cognitive functions (e.g., working memory).

### Connectome Integration and Segregation Significantly Mediates Age Effects on Maturation of the Primary-to-Transmodal Gradient

Functional integration and segregation (39), which correspond respectively to global information integrity and local information specialization within the brain network, have already emerged in neonates (40, 41) by the third trimester (40), and exhibit substantial changes throughout the developmental process (13, 14, 20). However, the primary-to-transmodal gradient is not observed at birth (8). Moreover, this gradient does not become the principal connectome gradient until early adolescence, as shown in this study. Thus, we investigate whether the integration and segregation of the functional brain network facilitates developmental changes in the core connectome gradient. To do this, we conducted a mediation analysis to examine the relationship between age, graph-theoretical metrics (which reflect network integrity and segregation), and global gradient measures (*SI Appendix*, Section 7-8).

We first examined the relationship between age and PC1 of the global gradient measures. Using a mixed linear model and controlling for gender and head motion parameters, we found that PC1 score increased with age (*t* = 5.33, *P* < 0.001, Fig. 4A). To evaluate network integration and segregation at the whole-brain and subsystem levels, we adopted the characteristic path length (*Lp*) and the clustering coefficient (*Cp*) of whole brain network (sparsity = 10%, unweighted) (42), and the modular segregation index (MSI) of each subnetwork (43), respectively, as the relevant variables in our analysis. For a given subnetwork (34), a positive MSI indicates greater segregation while a negative MSI indicates greater integration (43). We observed that the *Lp* and the *Cp*, as well as the MSI of the DMN, SMN, FPN and VN were positively correlated with age and PC1 (all *P* < 0.001, FDR-corrected *q* < 0.05, Fig. 4B and *SI Appendix*, Fig. S8). Using a non-parametric bootstrapping approach (10,000 times), we performed a mediation analysis in which age was taken as the independent variable, *Lp, Cp*, and the MSI of subnetworks were taken as the mediators, and PC1 of the global gradient measures was taken as the dependent variable (Fig. 4C). We found that the *Lp, Cp*, and MSI of the DMN, SMN, FPN and VN partially mediated the relationship between age and PC1 of the global gradient measures (indirect effect *ab*: *β* = 0.11, CI = [0.03, 0.19]; *β* = 0.13, CI = [0.05, 0.22]; *β* = 0.09, CI = [0.03, 0.16]; *β* = 0.09, CI = [0.04, 0.15]; *β* = 0.05, CI = [0.02, 0.09]; *β* = 0.03, CI = [0.01, 0.07], all bootstrapped *P* < 0.01, Fig. 4D and *SI Appendix*, Table S3). We also observed the mediation effects of the graph-theoretical metrics on age-associated changes in each global gradient measure (Fig. 4D).

**Fig. 4.**
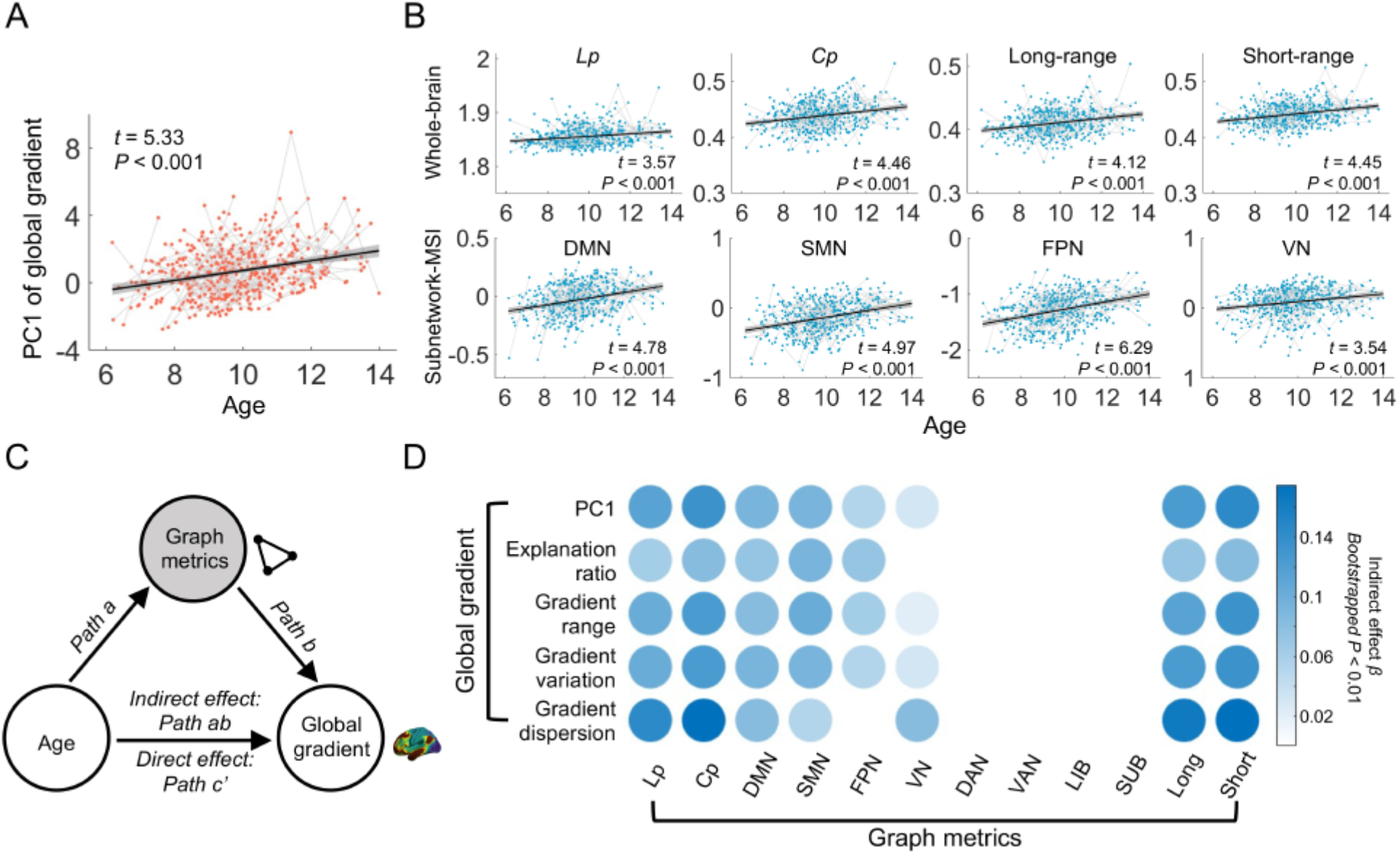
Mediation effect of integration and segregation in the functional brain network on age-associated changes in global gradient topography. Significant increases with age were observed in (**A**) PC1 of the global gradient measures and (**B**) multiple graph metrics, including *Lp, Cp*, the MSI of the DMN, SMN, FPN and VN, and the mean functional connectivity values of long and short range connections. (**C**) The mediation model, in which age was taken as the independent variable, graph metrics were taken as the mediator, and global gradient measures were taken as the dependent variable. Gender and head-motion parameters were included as covariates. (**D**) The graph metrics, including *Lp, Cp*, the MSI of the DMN, SMN, FPN and VN, and the mean values of long and short range connections significantly mediated age-associated changes in PC1 of the global gradient measure, as well as each individual global gradient measure. Most models showed a partial mediation effect, with the exception that the development of gradient dispersion was fully mediated by *Lp, Cp*, and functional connectivity values of long and short range connections. Gender and head-motion parameters were included as covariates. The color of circle represents the beta values of the indirect effect, and darker colors indicate a greater mediation effect. We assessed the statistical significance of the mediation effect using 99% bootstrapped (10,000 times) CIs. *Lp*, shortest path length; *Cp*, clustering coefficient; Long-range/Long, mean functional connectivity value of long range connections; Short-range/Short, mean functional connectivity value of short range connections; MSI, modular segregation index; DMN, default-mode network; SMN, sensorimotor network; FPN, frontoparietal network; VN, visual network; DAN, dorsal attention network; VAN, ventral attention network; LIB, limbic network; SUB, sub-cortical regions.

Considering that development in long and short range connections is a critical feature of functional brain networks in children (13, 44), we further investigated the influence of changes in connections of specific length on the development of gradient topography. Specifically, we calculated mean functional connectivity values for long (> 70 mm) and short (< 70 mm) range connections in each individual functional network (20). The mean functional connectivity values of both long and short range connections were positively correlated with age and PC1 (both *P* < 0.001, Fig. 4B and *SI Appendix*, Fig. S8). Using a mediation analysis (Fig. 4C), we found that the mean functional connectivity values of long and short range connections also partially mediated the relationship between age and global gradient topography (indirect effect *ab*: *β* = 0.12, CI = [0.06,0.20]; *β* = 0.14, CI = [0.07,0.22], all bootstrapped *P* < 0.01, Fig. 4D and Table S3). We also repeated the mediation analysis using each individual global gradient measure (Fig. 4D). Together, these findings suggest that the integration and segregation of the whole-brain network and specific subnetworks (i.e., the DMN, SMN, FPN and VN), as well as the strengthening of short and long range connections, promoted the maturation of connectome gradients in the developing brain.

### Maturation of the Primary-to-Transmodal Gradient is Associated with Gene Expression Profiles

Having documented the development of the two core connectome gradients during childhood and adolescence, here, we sought to explore whether changes in gradient property, as a manifestation of maturing hierarchical organization in the connectome, were linked to gene transcription profiles (for details, see *SI Appendix*, Section 9-10).

To this end, we examined the association between the rates of annual changes in regional gradient scores in the primary-to-transmodal gradient (i.e., as represented in the *β*-map, Fig. 5A), obtained from the results of our quantitative mixed linear model analysis, and the gene expression profiles from the Allen Human Brain Atlas datasets. First, we preprocessed the micro-array data according to a recommended pipeline (45) and obtained group-level gene expression maps that were normalized across samples and donors (10,027 genes). As the gene expression maps did not cover the whole brain at a voxel level, we matched both the gene expression maps and the *β*-map to a cortical parcellation atlas (46) (Fig. 5B). Next, we estimated the relationship between the *β*-map and each gene expression map using Pearson’s correlation (Fig. 5C). The significance of these correlations was obtained by permutation tests (10,000 times) in which spatial autocorrelations were preserved in the surrogate *β*-maps (36). We then ranked the genes according to the significance levels of their correlation coefficients. That is, genes were ranked in descending order, beginning with the genes with the most significant positive correlations and ending with the genes with the most significant negative correlations. Finally, we performed an enrichment analysis using the Gene Ontology enrichment analysis and visualization tool (GOrilla, http://cbl-gorilla.cs.technion.ac.il) (47). The significance of the GO enrichment terms were further tested using a null model, which was generated by performing an enrichment analysis using surrogate maps which preserved the empirically observed spatial autocorrelations (36). We found that genes within the gene set which showed significant enrichment were those involved in the biological processes of calcium ion regulated exocytosis and chemical synaptic transmission, as well as those associated with cellular components, specially, with axons and synapse parts (FDR-corrected, all *q* < 0.05, Fig. 5D, see *SI Appendix*, Table S4 for specific GO terms). We also performed Gene Ontology enrichment analysis on a gene set made up of the genes ranked in reverse order (i.e., starting with the most significant negative correlation and ending with the most significant positive correlation), but did not find significant enrichment of any genes in this set.

**Fig. 5.**
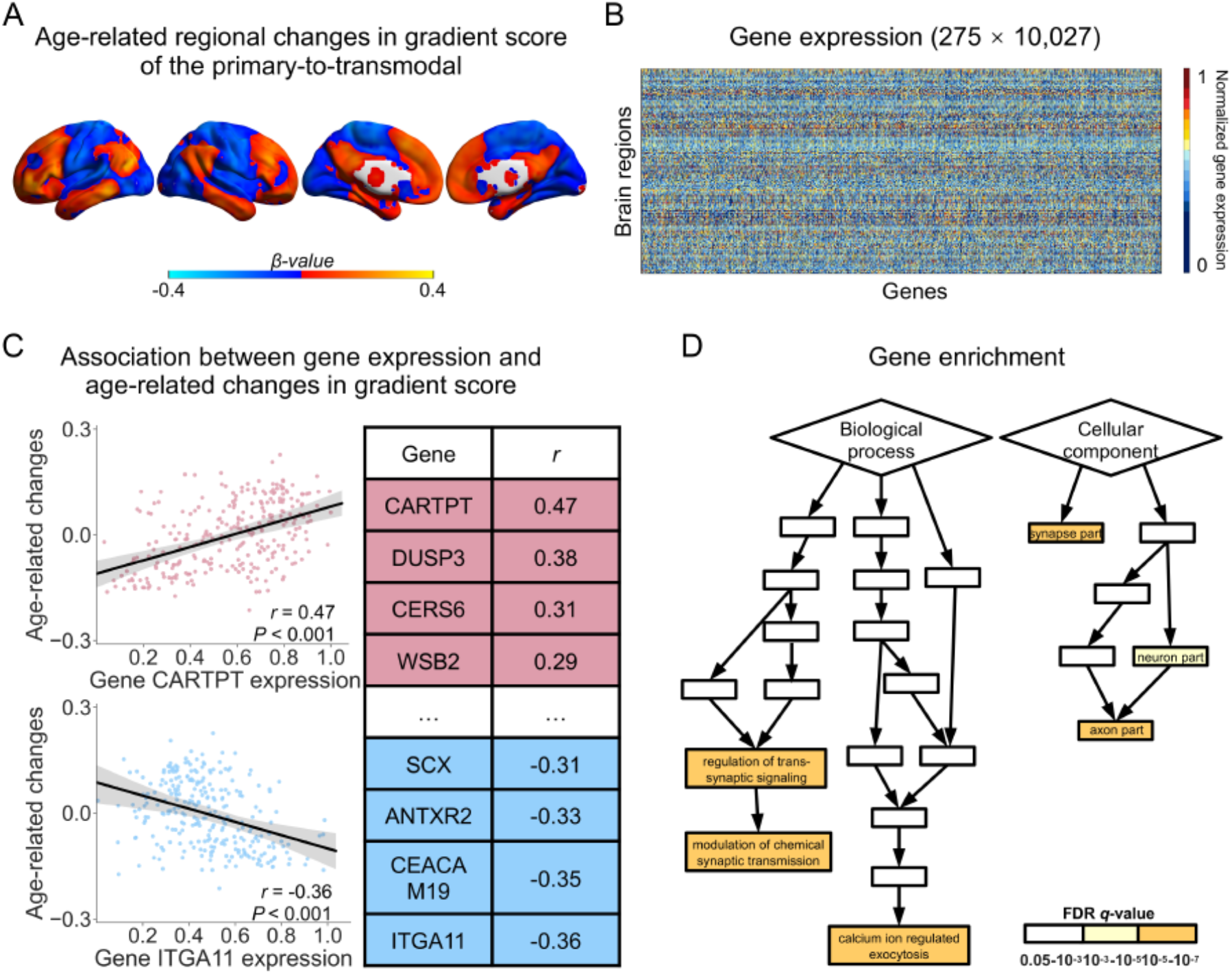
Association between maturation of the primary-to-transmodal gradient and gene expression levels. (**A**) The rates of annual change (age-related) in regional gradient scores (i.e., *β*-map) of the primary-to-transmodal gradient. (**B**) Normalized gene expression levels in 275 brain regions. (**C**) We calculated the correlation between the expression profile of each gene and the age-related *β*-map, and computed the statistical significance of the correlation using permutation tests (10,000 times) and generative modeling (36). Two representative scatter plots (both *P* < 0.001) were illustrated, with positive/negative correlations shown in red/blue. (**D**) Enrichment analysis. The gene set showed significant enrichment of genes associated with calcium ion regulated exocytosis and chemical synaptic transmission, and of genes related to axons and synapse parts (FDR-corrected, all *q* < 0.05).

### Validation Analyses

To demonstrate the robustness of our main findings, we performed several validation analyses (for details, see *SI Appendix*, Section 11), including: (*i*) estimating the longitudinal changes in connectome gradient properties using rs-fMRI data on which no global signal regression was performed during preprocessing (*SI Appendix*, Fig. S9); (*ii*) evaluating the influence of the sliding window method on observations of the emergence of the primary-to-transmodal gradient as the principal connectome gradient, this was undertaken by using only the last scan of each child and using different parameter combinations (window length = 20, step size = 5; window length = 30, step size = 10) (*SI Appendix*, Fig. S10); and (*iii*) assessing the mediation effect of graph-theoretical measures on developmental changes in global gradient topography, using brain networks with different sparsities (i.e., 5% and 15%) (*SI Appendix*, Fig. S11 and S12). Overall, the application of different strategies for analysis did not affect or alter our main conclusions.

## Discussion

The present longitudinal study documents the developmental stages and related gene expression profiles of the primary-to-transmodal gradient in the functional connectome. Specifically, the primary-to-transmodal gradient exhibits a highly dynamic nature from childhood to adolescence, with longitudinal changes including emergence as the principal connectivity gradient and expansions of global topography, as well as focal tunings in the primary and transmodal regions. Moreover, the maturation of this gradient is mediated by changes in functional brain network integration and segregation, and is associated with improvement in the performance of working memory as well as the expression levels of genes involved in calcium ion regulated exocytosis, synaptic transmission, axons, and synapse parts. These results shed new light on the stages in the maturation of and the biological basis behind the core connectome hierarchy, and also have implications for understanding normal cognitive development and neuropsychiatric disorders related to abnormal development.

Prior studies on the adult brain have demonstrated on the existence and topography of the principal primary-to-transmodal gradient in the human brain functional connectome (6). This principal gradient characterizes a spectrum ranging from perception and action to abstract cognitive functions. Recently, three developmental and lifespan studies on functional connectome gradients have been also been conducted. Larivière et al. (8) investigated functional gradients in neonates, and found that the primary-to-transmodal gradient was not fully developed at birth. Bethlehem et al. (7) used cross-sectional rs-fMRI data to study age effects on the primary-to-transmodal gradient between the ages of 18 to 88 years. Dong et al. (9) compared functional gradients in three predefined age groups (< 12 years old, 12-17 years, and adults) and showed that the primary-to-transmodal gradient became the dominant gradient in the second and third groups (12-17 years and adults). However, these three studies did not investigate when the core primary-to-transmodal gradient emerges or how it develops during childhood and adolescence. Thus, a longitudinal and quantitative delineation of the developmental trajectory for the primary-to-transmodal gradient is still missing from the literature. Moreover, the three above mentioned studies also did not examine how connectome gradient development is related to the maturation of specific cognitive functions and gene expression profiles.

Between childhood and adolescence, individuals undergo a transition from the concrete operational stage into the formal operational stage, requiring the formation of abstract conception and logical thinking (10, 49). Thus, the primary-to-transmodal gradient’s replacement of the sensorimotor-to-visual gradient as the principal functional connectivity gradient may be a consequence of the requirement to adapt to the rapid development of complex and abstract cognitive functions during this period. As an anchor of the primary-to-transmodal gradient, the DMN, which is involved in multiple high-order abstract cognitive functions (50), undergoes remarkable alterations over the same developmental period. These include sub-regional reorganization (16) and strengthened long range connections (13, 18, 51). Notably, Larivière et al. also speculated that the absence of adult-like principal gradient patterns in neonates (8) may be compatible with the underdeveloped DMN connectivity pattern observed at birth (41, 52, 53).

Interpreted in conjunction with previous studies, we speculated that the emergence and progressive dominance of the adult principal gradient may be attributed to the maturation of DMN, which also supports cognitive growth during childhood and adolescence. Moreover, the development of the primary-to-transmodal gradient involves the expansion of global topography and increases in spatial variation, as well as focal tunings in primary and transmodal regions. Previous studies have reported that compared to adults, children have a lower level of functional segregation between the DMN and other brain systems, as well as a lower level of functional integration between the anterior and posterior regions of the DMN (13, 17, 19, 51). These results, together with our findings, reflect an increasing segregation between the primary sensory network and the transmodal regions of the DMN with age. Such observations correspond well with the “tethering hypothesis”, which proposes that the increasing functional discrepancies between the transmodal and the primary areas facilitates the formation of abstract cognition by avoiding the interference of input noise (54). The maturation of abstract thinking capacities during childhood and adolescence are closely related to the development of working memory, which is a cognitive system that temporarily holds the information necessary for complex high-order cognitive functions (55, 56). A better working memory capacity allows to more items, along with their associations, to be stored in temporary processing space, which is essential for the formation of more complex thoughts (55, 56). Thus, the development of working memory enables a child to hold more items in their mind at a time, resulting in the maturation of abstract cognition. Here, our observation of a positive association between the expansion of global topography in the principal gradient and enhanced performance in individual working memory may indicate that the segregation of transmodal regions supports the development of abstract cognition.

The human brain functional connectome is able to achieve efficient global information communication at a low wiring cost. This capacity is heavily dependent on the optimized balance between segregation and integration (39, 57). The results from our longitudinal analysis of the relationship between age and network topology showed that, between childhood and adolescence, the functional connectome exhibits a continuous increase in segregation and reduction in integration, as reflected by the increases in clustering coefficient, characteristic path length, and subnetwork modular segregation index. However, in contrast to our results, several prior studies have reported non-significant age-related differences in the two network parameters in children (19, 58). These differences are partly due to the cross-sectional nature of the prior studies and the specific age groups selected for comparison. Nonetheless, our results remain supported by the findings and evidence from a large number of previous studies. Firstly, the formation of specific brain networks between childhood and adolescence is believed to be supported by both experience dependent evoked neuronal activity and spontaneous coordinated neuronal activity (14). Several rs-fMRI studies in children have also shown that as age increases, higher-order cognitive networks such as the DMN and the frontoparietal networks tend to become more functionally segregated from other brain systems (17, 18, 51, 58). The findings from these past studies align with our observations that the connectome exhibits enhanced functional segregation with increasing age. Secondly, microstructural studies using histological and postmortem data have revealed the existence of overproduced dendrites and axons during the perinatal phase, which leads to redundant synaptic connectivities in postnatal brain development (59, 60). These redundant connectivities are maintained throughout childhood, but are reduced to a large extent during adolescence due to synaptic pruning (61, 62). Several rs-fMRI studies have also shown that functional connectivity (e.g., subcortical-cortical) in the child brain is overabundant and more diffuse when compared to that in adolescent and adult brains (13, 58). These results suggest an overall reduction of functional integration levels with age. Importantly, in the present study we demonstrate that age effects on the global topographic profile of connectome gradients are mediated by the changes in segregation and integration of the functional networks. The dynamic reconfiguration of the global brain network may gradually form a hierarchical network structure, which can be efficiently linked to support information transfer between nodes at low wiring costs. Together, these findings highlight the significance of integration and segregation in the brain network in promoting the development of connectome hierarchy.

Our connectome-transcriptome association analysis established a link between longitudinal changes in the primary-to-transmodal gradient and the enriched expression of genes involved in the biological processes of calcium ion regulated exocytosis and chemical synaptic transmission. The chemical synaptic transmission process includes the release of neurotransmitter molecules from a pre-synapse, the transport of these molecules across a chemical synapse, and the subsequent activation of neurotransmitter receptors at the post-synapse (63). The calcium ion plays an important role in the release of neurotransmitters. Specifically, the activity and distribution of calcium ions in the presynaptic membrane area can regulate the timing and intensity of synaptic transmission (64). Impaired neurotransmitter release introduced by the abnormal activity and distribution of calcium ions is associated with neurodevelopmental disorders. Autism-associated mutations of the RIM3 gene can suppress calcium ion inactivation, leading to impaired neurotransmitter release in rodent model (65). Other prior studies have also suggested that synaptic transmission is associated with structural and functional development of the brain during childhood and adolescence (29, 66). For example, cortical thinning and intracortical myelination of the association cortex are spatially associated with the expression level of genes related to the synaptic transmission of glutamate (66). Similarly, prefrontal function and connectivity is influenced by synaptic transmission in dopamine signaling (67), which is also associated with individual variability in behavioral phenotypes (68).

In addition, the development of the primary-to-transmodal connectome gradient was also associated with genes related to cellular components, specifically those related to axons and synapse parts. During childhood and adolescence, the brain undergoes refinements in various aspects of its microstructural features (12), including the pruning of synaptic density (mainly in prefrontal cortex) (62) and the continuous myelination of axons (69). Synaptic pruning may eliminate redundant connections in the transmodal regions, while the myelination of axons may reflect the strengthening of connections involved in external experience and learning. Both synaptic pruning and axon myelination result in the rewiring of structural pathways, which is likely to lead to re-organization in the macroscale functional connectome during childhood and adolescence. Numerous studies have reported functional segregation between the primary cortex and the transmodal regions during this stage (15, 19, 51). Thus, the principal connectome gradient, anchored by the primary areas and the transmodal regions, may capture an overwhelming proportion of the connectivity changes resulting from synaptic pruning and axon myelination across the whole-brain connectome, and gradually becomes the dominant gradient in late childhood. Using diffusion MRI data in children, several studies have demonstrated that the fractional anisotropy of white matter, which is an indirect indicator of axon myelination, is correlated with the development of working memory (70). This corresponds with our observation of a direct connection between primary-to-transmodal gradient development and enhancements in working memory. Hence, our findings are compatible with previous studies, and provide evidence from a different perspective to support the link between the expression of axon and synapse part related genes, connectome gradient development, and the maturation of specific cognitive functions. Based on the above, it also seems plausible that the development of hierarchical architecture in the functional connectome is underpinned by calcium ion regulated exocytosis and chemical synaptic transmission processes acting together with dynamic changes in axon and synaptic structures.

Several issues need to be considered. Firstly, previous work has suggested that the functional connectome can be anatomically shaped by the structural connectome (71) and that the relationship between the structural and functional connectomes can be modulated by age (72). A recent study also reported that spatial variability in structure-function connectivity coupling is aligned with cortical hierarchies during youth (73). The current study delineates the stages in the maturation of the primary-to-transmodal gradient in the functional connectome, and notes that this process involves changes in both global and regional topography. However, how gradient development in the child functional connectome may be constrained or shaped by the structural connectome remains to be further elaborated. Secondly, gene expression maps used in this study were constructed using data obtained from postmortem adult brains, and gene expression levels may vary between adults and children (74). A recent study has evidenced that the spatial topology of gene expression in the visual cortex undergoes continuous development before the age of 5 (75). Our findings in respect of the relationship between the development of the primary-to-transmodal gradient and gene transcriptome profiles should be interpreted cautiously. Nonetheless, the gene expression maps from the Allen Institute for Brain Science are the most spatially detailed dataset available with standard coordinates provided. In future, the availability of a pediatric specific gene expression dataset will provide stronger and more conclusive evidence on this issue. Finally, the developmental profile of the primary-to-transmodal gradient described in this study is that observed in typically developing children. A recent study revealed that the primary-to-transmodal connectome gradient shows a contracted pattern in children with autism (32), providing a new perspective for understanding the interplay of low and high-level cognitive deficits. Further studies should be conducted to explore whether disruptions in the connectome gradient can serve as biomarkers for the diagnosis and treatment evaluation of neuropsychiatric disorders related to abnormal development (e.g., autism and attention-deficit/hyperactivity disorder).

## Materials and Methods

### Brain Imaging Dataset

We collected a longitudinal rs-fMRI dataset from 305 children (Discovery Dataset, 491 scans taken at three points in time from 162 males and 143 females aged 6-14) and a cross-sectional sample of 61 adults (ages 18-28, 25 males and 36 females), as a part of the Children School Functions and Brain Development Project in China (Beijing Cohort). All individuals were cognitively normal and had no history of neurological/psychiatric disorders, substance abuse (including illicit drugs and alcohol), significant head injury, physical illness, or MRI contraindications. The study was approved by the Ethics Committee of Beijing Normal University, and written informed consent was obtained from the participants or their parents/guardians prior to commencement of the study. We also used a Replication Dataset to validate our main results, which was downloaded from the NKI-RS publicly shared website (http://fcon_1000.projects.nitrc.org/indi/enhanced/). In this Replication Dataset, rs-fMRI data was collected from 254 children (390 scans taken at three points in time from 145 males and 109 females aged 7-18). All the children were cognitively normal and had no mental or neurological disorders.

### Behavioral Data

The cognitive performance of participants included in the Discovery Dataset were assessed using a classic numerical *N*-back working memory task and a child-friendly version of the Attention Network Task (ANT) implemented during task-state fMRI (76). We collected 454 scans of individual performance in the N-back working memory task, and calculated the 2-back d-prime (i.e., the inverse of the cumulative Gaussian distribution of the 2-back hit ratio subtracted by the inverse of the cumulative Gaussian distribution of the 2-back false alarm ratio (77)) of 409 scans (ages 6-14, 194 males and 215 females) after data cleaning (exclusion criteria: 0-back hit ratio < 0.5 or 1-back hit ratio = 0 or 2-back hit ratio = 0). In addition, we also assessed 418 scans (after data cleaning) of ANT performance (ages 6-14, 202 males and 216 females), measuring mean response times for altering, orienting and execution attention. Specific data cleaning criteria can be found in Hao et al. (76).

### Gene Expression Dataset

Microarray data from six donors (mean age: 42.5 y, 5 males and 1 female) was downloaded from the Allen Human Brain Atlas website (http://human.brain-map.org, RRID: SCR_007416). The six adult brains were parceled into 3,702 spatially distinct tissue samples, with each tissue sample capable of being mapped back into Montreal Neurological Institute coordinate space, and the expression levels of more than 20,000 genes were quantified (27). Gene expression level was normalized across samples within a single brain, and across donors to minimize non-biological confounding factors while maintaining biologically relevant variance.

### Data Analysis

Using the preprocessed rs-fMRI images and gene expression profiles, we performed the following analyses:

i. To characterize how core connectome hierarchy develops during childhood and adolescence, we conducted a gradient decomposition analysis (6) on the voxel-wise functional connectivity matrix of each individual, and performed both a qualitative cross-participant sliding window analysis and a quantitative mixed linear model analysis to examine the longitudinal changes in global topography and regional gradient scores of functional connectivity gradients in the connectome.
ii. To examine the cognitive implications of connectome gradient development, we charted the age-associated *t*-statistical map of the connectome gradients against cognitive activity maps from the Neurosynth dataset (https://neurosynth.org/) (35) and examined the direct link between individual primary-to-transmodal gradient measures to performance of working memory and attentional ability.
iii. To assess whether and how the integration and segregation of functional brain networks facilitates the maturation of the primary-to-transmodal gradient, we performed a mediation analysis to examine the relationship between age, graph-theoretical metrics (i.e., *Lp, Cp*, subnetwork MSI, and the mean functional connectivity values of long and short range connections), and the global connectome gradient measures.
iv. To explore potential gene expression profiles that are associated with the maturation of the primary-to-transmodal gradient, we conducted a Pearson’s correlation analysis followed by a Gene Ontology enrichment analysis.
v. To validate our main findings, we re-analyzed the Discovery Dataset using a number of different approaches, including re-analyzing without performing global signal regression during rs-fMRI data preprocessing, re-performing the sliding window analysis using cross-sectional data and different parameters, and re-performing analyses involving graph-theoretical metrics using different specifications of network sparsity.

See the *SI Materials and Methods* section of the *SI Appendix* for further details.

## Acknowledgements

The study was supported by the National Natural Science Foundation of China (Nos. 31830034, 82021004, 81620108016, 31221003, 31521063, 81671767, 82071998, and 81971690), Changjiang Scholar Professorship Award (No. T2015027), National Key Research and Development Project (No. 2018YFA0701402), Beijing Nova Program (No. Z191100001119023), the Beijing Brain Initiative of Beijing Municipal Science & Technology Commission (No. Z181100001518003), and the Fundamental Research Funds for the Central Universities (No. 2020NTST29). We thank the National Center for Protein Sciences at Peking University in Beijing, China, for assistance with MRI data acquisition. We also thank the Allen Institute for Brain Science for providing the gene expression data.

## Author contributions

Conceptualization: Y.X., M.X., Y.H.; Data collection: W.M., Y.W., Z.P., J.Luo, S.P., M.C., L.H., S.Tan, J.G., S.Q., S.Tao, Q.D.; Methodology: Y.X., M.X., J.Liu, X.Liao, T.L., X.Liang, T.Z., Z.S., L.S., G.G.; Data analysis: Y.X., M.X.; Supervision: Y.H.; Writing—original draft: Y.X., M.X., Y.H.; Writing—review & editing: Y.X., M.X., Y.H.

## Competing interests

Authors declare that they have no competing interests.

## Data and materials availability

All data are available in the main text or the supplementary materials.

## Supporting Information (SI) Appendix

SI Materials and Methods

Fig. S1. The five most significant connectome gradients in the adult brain.

Fig. S2. The range of the age windows in children.

Fig. S3. Longitudinal changes in the primary-to-transmodal gradient, with gender, head-motion parameters, and the sensorimotor-to-visual gradient included as covariates.

Fig. S4. Longitudinal changes in the sensorimotor-to-visual gradient, with gender and head-motion parameters included as covariates.

Fig. S5. Longitudinal changes in the core connectome gradients in the Replication Dataset.

Fig. S6. Statistical group comparisons (children vs. adults) of the two connectome gradients.

Fig. S7. Correlation between global measures of the primary-to-transmodal gradient and performance in the working memory task (2-back d-prime).

Fig. S8. Correlation between graph metrics of the functional brain network and global topography of the primary-to-transmodal gradient.

Fig. S9. Longitudinal changes in the two connectome gradients, as observed in rs-fMRI data with no global signal regression, and with gender and head-motion parameters included as covariates.

Fig. S10. Explanation ratio of the two gradients across age windows.

Fig. S11. Mediation effect of integration and segregation in the functional brain network (sparsity=5%) on age-associated changes in global gradient topography.

Fig. S12. Mediation effect of integration and segregation in the functional brain network (sparsity=15%) on age-associated changes in global gradient topography.

Fig. S13. Longitudinal changes in dispersion within and between subnetworks.

Table S1. Brain regions showing significant age-related changes in gradient score.

Table S2. Brain regions showing significant correlation between gradient score and working memory performance.

Table S3. Graph metrics showing significant mediation effect on age-associated changes in the PC1 score of global measures of the primary-to-transmodal gradient.

Table S4. Gene Ontology enrichment analysis of genes associated with development of the primary-to-transmodal gradient.

Movie S1. Developmental dynamics of functional connectome gradients across different ages.

## Notes

### Competing Interest Statement

The authors have declared no competing interest.

